# Prefrontal cortex proteomic alteration after social instability stress in adolescents rats

**DOI:** 10.1101/2023.05.08.539806

**Authors:** Evelyn C. S. Santos, Ana Filipa Terceiro, Rui Vitorino, Igor Lopes, Renata L. Alves, João B. Relvas, Teresa Summavielle, Ana Magalhães

**Affiliations:** Institute of Research and Innovation in Health (i3S) and Institute for Molecular and Cell Biology (IBMC), University of Porto, Porto, Portugal, Glial Cell Laboratory; Institute of Research and Innovation in Health (i3S) and Institute for Molecular and Cell Biology (IBMC), University of Porto, Porto, Portugal, Addiction Biology Laboratory; iBiMED-Department of Medical Sciences, Institute of Biomedicine, University of Aveiro, Aveiro, Portugal; Faculty of Medicine of the University of Porto (FMUP), Porto, Portugal

**Author notes:** Equally contributed to this work (Evelyn C. S. Santos, Ana Filipa Terceiro).

## Abstract

Early life stress can have significant effects on the developing brain and lead to changes in the structure and function of brain regions involved in stress regulation, emotion and cognitive control. Here, we used the social instability stress (SIS) protocol to understand the impact of social stress during mild (PND30) and late (PND45) adolescence. Our results revealed that SIS can compromise the dominance-subordination coping strategy but does not affect social recognition and motivation in rats. Moreover, SIS can lead to subtle modifications at the molecular level that hamper normal development of the prefrontal cortex in a sex- and age-dependent manner. Understanding the impact of early life stress on brain organization is crucial for developing effective prevention and intervention strategies. By identifying those who are most vulnerable to the effects of stress and providing targeted support and resources, it may be possible to mitigate the negative consequences of early adversity and promote healthy brain development.

## Introduction

It is widely known that stressful events affect neurobehavioral responses (Blanchard *et al*., 2001; Schneiderman *et al*., 2005; Sandi & Haller, 2015). Early life stress can have profound and lasting effects on brain organization and has been strongly associated with several psychopathologies (McEwen, 2000), with lasting effects into the life of adult individuals (Smith *et al*., 2018).

Stressors can be described and divided into several different categories. Among them, social stress is associated with the development of negative effects on mental health and well-being. We and others have previously reported that early social stress, such as maternal separation, leads to substantial negative effects on brain function and behavior in rodents (Makinodan *et al*., 2012; Carlyle *et al*., 2012; Nishi, 2020; Bonnefil *et al*., 2019 Alves *et al*., 2022). Research shows that social stress is particularly damaging during adolescence, a critical period of functional and structural development. Behaviorally, adolescents are more sociable, form hierarchical peer relationships, and are more sensitive to acceptance and rejection by their peers than younger individuals (Blakemore, 2012); but they are also impulsive, active seekers of new sensations and willing to take more risks (Crews & Hodge, 2007). Social changes occurring during adolescence contribute to the development of the social brain, as demonstrated by Blakemore and colleagues. Using neuroimaging studies, they showed that there are higher levels of activity in the prefrontal cortex in adolescence as compared to childhood or adulthood (Blakemore, 2008).

The prefrontal cortex (PFC) is particularly critical in the adolescence period. It is a brain region vulnerable to developmental experiences and important to establish the final organizational phase of neural maturation during adolescence (Semeralul *et al*., 2006; Caballero *et al*., 2016; Uytun, 2018). Furthermore, the PFC has been implicated in the regulation of emotion, social cognition, and decision-making. Impairments in executive function in the PFC are symptoms of several stress-related psychological disorders (e.g., schizophrenia, depression, PTSD) (Cerqueira *et al*., 2007; Arnsten, 2009; Arnsten *et al*., 2015; Riboni & Belzung 2017). Therefore, understanding aspects that influence the brain maturation, particularly in the PFC, and remain under construction during adolescence is crucial (Arain *et al*., 2013).

In this context, it is important to identify the mechanisms by which social stress alters the developing brain and to identify resilience and risk factors to stress during adolescence. However, there has been little research into the impact of social stress during different periods of adolescence despite late adolescence being a period of significant social development.

In preclinical models, social stress can be assessed by different tests, such as the Social instability stress (SIS) (for review McCormick *et al*., 2021). The SIS consists of unstable social hierarchical dynamics for several days or weeks to induce negative valence behaviors and physiological impairments (Yohn *et al*., 2019). The unstable social dynamics are obtained by unpredictability and social stress based on daily rotation of the cage group composition, thereby exposing animals to unfamiliar same-sex cage partners regularly (for review Koert *et al*., 2021). This is an effective paradigm in both male and female rodents and an etiologically relevant preclinical model of social stress (Yohn *et al*., 2019; Koert *et al*., 2021) and has been proposed as a valid model for social disorganization in urban areas (Tabibzadeh & Liisberg, 1997).

The effects of SIS on social behavior during the adolescent period were explored by Hodges and colleagues (2017; 2018, 2019) and they found that social instability stress impaired social recognition and increased the time in social approach but reduced the time in social interaction with an unfamiliar peer (Hodges *et al*., 2017; 2018). However, both SIS and control rats did not differ in how they found social interactions with unfamiliar peers rewarding in a conditioned place preference task (Hodges *et al*., 2017).

The current study uses the SIS paradigm to investigate the social behavioral and molecular adaptations in mild and late phases of adolescence. To evaluate social behavior, we assess social motivation, social memory/recognition, and dominance. These behavioral domains can be conceptually compared to the human classifications: social motivation, knowledge of self and others, and hierarchies within groups (Bicks *et al*., 2015). We focus on the PFC due to its sensitivity to developmental experiences, suggesting that it is a critical region to consider in the perspective of adolescent brain maturation. Determining how social stress influences PFC functioning in males and females at different developmental stages will deepen our knowledge of brain-behavior relationships and the mechanisms underlying sex-biased disorders.

## Materials and methods

### Animals

Sprague Dawley rats originally obtained from Charles River (Barcelona/Spain) were used in this study. All animals were kept under standard conditions (21±1°C; 60±5% humidity; 12h light/12h dark cycle - light off at 12h00) with ad libitum access to food and water. Mating was done by introducing a male into a cage with two females (nulliparous) at the beginning of the dark cycle. Pregnant rats were then housed individually from gestational day 16 until delivery, designated as postnatal day (PND) 0. On PND 1, all litters were sexed balanced (4/5 females and 4/5 males). Rats were weaned on PND 21 and randomly assigned to the following experimental groups: 1) SIS-PND30 - Social instability experienced during mild adolescence period (from PND 30 to 37); 2) SIS-PND45 - Social instability experienced during the late adolescence period (from PND 45 to 52); and no cage mate alterations control groups: 3) Control-PND30 and 4) Control-PND45.

Experiments involving animals were approved by the Portuguese regulatory agency Direcção Geral de Alimentação e Veterinária (DGAV) and the animal ethics committee of IBMC-i3S. The animal facility and the people directly involved in animal experimentation were also certified by DGAV. All animal experiments considered the Russell and Burch 3R’s principle and followed the European guidelines for animal welfare (2010/63/ EU Directive).

### Social instability stress model

The social instability stress model (SIS) was adapted from McCormick and colleagues (2015) with some modifications. Animals were paired with a new cage partner of the same age and sex, also experiencing the SIS procedure (Fig 1). The pairing with a new cage partner occurs every 12h, for 7 days. On day 8, all animals returned to their original cage partner, ending the SIS experience. Rats were then left undisturbed except once a week for cage maintenance until behavioral tests. Control animals were only disturbed once a week for cage maintenance except during the SIS procedure period during which the cages were removed from the rack and placed back so that the number of times the cages were moved was the same as in the experimental group.

**Figure 1.**
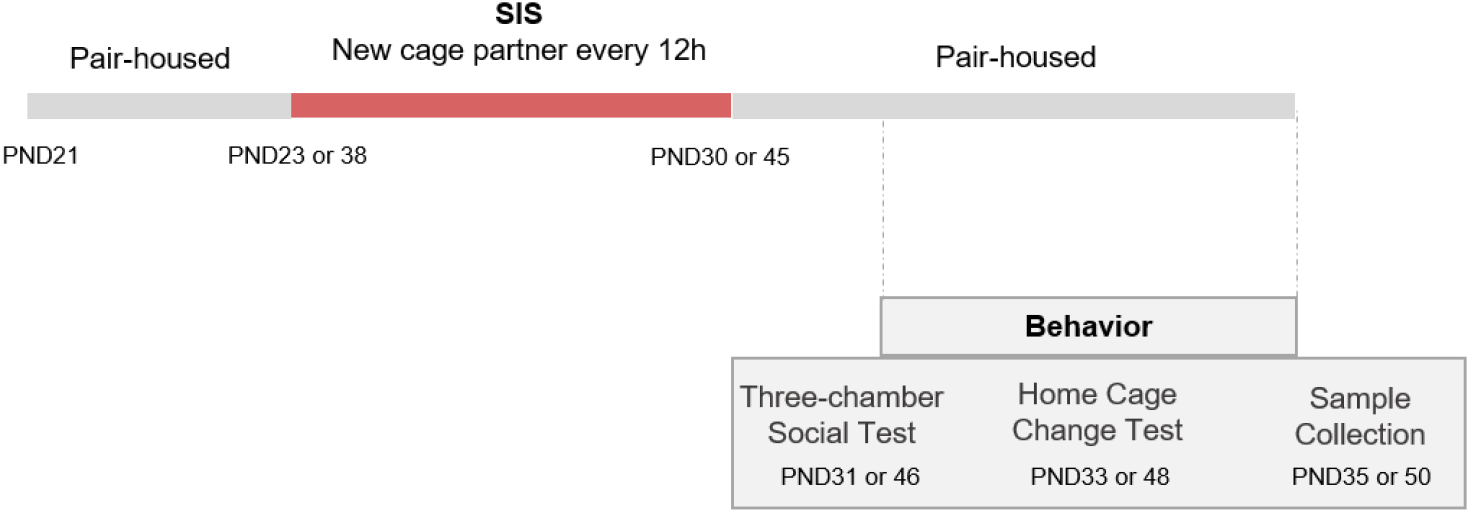
Schematic diagram of experimental protocol. Animals were pair-housed after weaning and submitted to social instability stress (SIS) protocol. Social behavioral analysis was conducted after the SIS in two distinct aged rats, mild (PND 30) and late (PND 45) adolescence.

This procedure differs from McCormick and colleagues (2015) in three main points: 1) animals were not previously isolated and restrained in small containers for 1 hour; 2) The procedure lasted 7 days instead of 15 days; and 3) animals have two new partners and cages each day, instead of 1 new partner and cage each day. However, in the two protocols, all rats interact with 15 different peers.

### Behavioural Tests

To evaluate the impact of SIS during two periods of development on social cognition, the following tests were performed: 1) Three Chamber Social Test, to evaluate social motivation and social recognition (Silverman *et al*, 2010), and 2) Novel Cage Test, to assess perceptive dominance/subordination relationships in a home-cage (Magara *et al*., 2015). All time measurements were recorded in seconds and distances in centimeters.

### Three chamber sociability and social novelty preference tests

The three-chamber test evaluates the rat preference in two different social paradigms: 1) Sociability test, used to evaluate social motivation and is measured by the amount of time spent with another rat, as compared to time spent alone in an identical but empty chamber (Fig. 2A); 2) Social preference test, used to evaluate components of social affiliation, social recognition, and social memory and, is measured by the amount of time spent with an unfamiliar (novel) rat rather than with a familiar one (Fig. 2 D) (Silverman *et al*, 2010).

**Figure 2.**
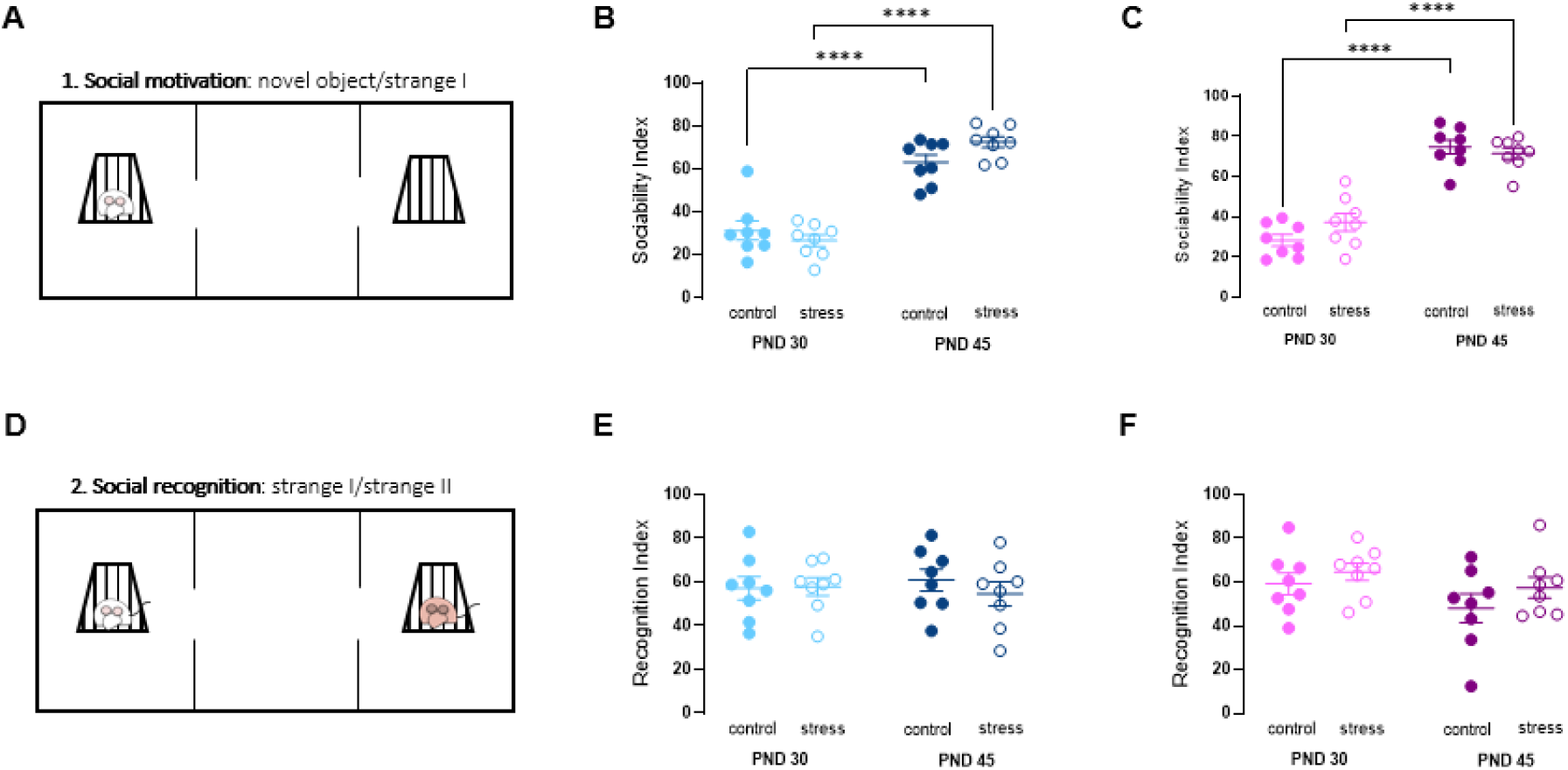
Effect of SIS on mild (PND 30) and late (PND 45) adolescence in the Three Chamber Sociability test. (**A**) Schematic representation of the Social motivation session; Sociability index of mild and late male (**B**) and female (**C**) adolescents. (**D**) Schematic representation of Social recognition session; Recognition index of mild and late male (**E**) and female (**F**) adolescents. Two-way ANOVA with Sidak’
ss comparisons. *n*= 8 per group. Data are expressed as mean ± SEM. \*\*\**p* < 0.001.

The apparatus consists of a box divided into three connected rectangular chambers (40.5 × 80 × 40 cm). From the central compartment, two openings provide access to each side compartment. The central compartment was the starting point for each test. The rat was allowed to roam around the three chambers for 10 minutes as a habituation period. Then, the sociability test started by putting a wire box in the middle of each side compartment. One of the wire cups acts as the novel object (empty) and as the container control for the novel object, whereas the other cup that encloses the stranger rat serves as a social stimulus. The widely spaced wire bars of the wire cup permit olfactory, visual, auditory and some tactile contact while preventing other kinds of social interactions, thus ensuring a measure of interest in approaching and remaining in physical proximity to one another (Silverman *et al*., 2010). The sociability test is initiated by placing the experimental rat in the central chamber and allowing the exploration of the apparatus for 10 minutes. The experimental rat returns to the home cage while the new rat (unfamiliar social stimulus) was added to the empty wire box for the social novelty preference test. Here, the previous rat (social stimulus) becomes the familiar rat. The experimental rat was then put back in the central chamber and was allowed to explore the 3 chambers for an additional 10 minutes. After each trial, the chambers and wire cages were cleaned with neutral, odor free soap. The time spent in contact with each wire box and compartment was measured using the software The Observer XT v7.0, Noldus, Netherlands. The sociability index was estimated by measuring the difference in time spent by the rat between the empty and social wire box compartment divided by the sum of the time spent exploring both wire boxes. The recognition index is calculated by measuring the difference in time spent by the rat between the familiar and unfamiliar rats divided by the sum of the time spent exploring both rats.

### Home Cage Change Test

On PND 39/54, dominant behavior was assessed by performing the Home Cage test according to the protocol described by Magara *et al* (2015). Two house-paired rats were moved into a new home cage (Type III) with new bedding material to simulate a cage change. Their behavior was video-recorded for 10 min under red light of about 3 lux intensity. The rats were marked with a pen on their back for identification. The duration and frequency of behaviors were analyzed per Magara and colleagues (2015) description (Supplementary Table 1). The analysis was conducted manually with the event-logging behavior quantification software BORIS (Behavioral Observation Research Interactive Software, Torino, Italy (Friard & Gamba 2016). Behaviors were grouped in 4 categories (neutral, dominant, submissive and aggressive) and the percentage of each behavioral category was calculated.

### Statistical analyses

Comparisons between groups were done by a two-way ANOVA with *Age* (PND30, PND 45) and *condition* (control, peer stress) as independent factors. When some of the factors (age, treatment or *sex*) was found not significant (*p* > 0.05), it was removed, data were collapsed across, and a two-way ANOVA was conducted. Sidak’
ss *post-hoc* test was used to perform multiple comparisons. Statistical significance is represented in this article by the following symbols: * *p* ≤ 0.05, ** *p* ≤ 0.01, and *** *p* ≤ 0.001. All data were analysed with GraphPad Prism v. 8 (GraphPad Software Inc.).

### Proteomic analysis

#### Protein extraction, fractionation and digestion

On PND35 for mild adolescent rats and PND50 for the late adolescence group, 2 days after the last behavioural test, rats were anesthetized, transcardially perfused with PBS and the PFC was collected and stored at -80ºC. Tissues were lysed using RIPA buffer (50 mM Tris, 150 mM NaCl, 1% Triton X-100, 0.5% Sodium Deoxycholase, 0.1% SDS) supplemented with a cocktail of phosphatase inhibitors (50 mM NaF, 1.5 mM Na_3_VO_4_) and protease inhibitors (0.1 mM PMSF, 1 mM CLAP (chymostatin, lemptpin, antipapin and pepstatin A) and 1 mM DTT). Samples were sonicated, rested on ice for 30 min and centrifuged at 16 000 rpm for 10 min (4ºC). The supernatants were collected, and protein concentration was determined using the BCA Protein Assay Kit (Pierce, Thermo Scientific), according to the manufacturer’s instructions.

#### Proteomics Data Acquisition

Proteins were solubilized with 100 mM Tris pH 8.5, 1% sodium deoxycholate, 10 mM tris(2-carboxyethyl) phosphine (TCEP), 40 mM chloroacetamide and protease inhibitors for 10 minutes at 95ºC at 1000 rpm (Thermomixer, Eppendorf). Each sample was processed for proteomics analysis following the solid-phase-enhanced sample-preparation (SP3) protocol as described in Huges and colleagues (2019). Enzymatic digestion was performed with Trypsin/LysC (2 micrograms) overnight at 37ºC at 1000 rpm.

Protein identification and quantitation was performed by nanoLC-MS/MS as previously described (Osorio *et al*., 2021). This equipment is composed by an Ultimate 3000 liquid chromatography system coupled to a Q-Exactive Hybrid Quadrupole-Orbitrap mass spectrometer (Thermo Scientific, Bremen, Germany). Samples were loaded onto a trapping cartridge (Acclaim PepMap C18 100Å, 5 mm x 300 μm i.d., 160454, Thermo Scientific) in a mobile phase of 2% ACN, 0.1% FA at 10 μL/min. After 3 min loading, the trap column was switched in-line to a 50 cm by 75μm inner diameter EASY-Spray column (ES803, PepMap RSLC, C18, 2 μm, Thermo Scientific, Bremen, Germany) at 250 nL/min. Separation was generated by mixing A: 0.1% FA, and B: 80% ACN, with the following gradient: 5 min (2.5% B to 10% B), 120 min (10% B to 30% B), 20 min (30% B to 50% B), 5 min (50% B to 99% B) and 10 min (hold 99% B). Subsequently, the column was equilibrated with 2.5% B for 17 min. Data acquisition was controlled by Xcalibur 4.0 and Tune 2.11 software (Thermo Scientific, Bremen, Germany).

The mass spectrometer was operated in data-dependent (dd) positive acquisition mode alternating between a full scan (*m/z* 380-1580) and subsequent HCD MS/MS of the 10 most intense peaks from full scan (normalized collision energy of 27%). ESI spray voltage was 1.9 kV. Global settings: use lock masses best (*m/z* 445.12003), lock mass injection Full MS, chrom. peak width (FWHM) 15s. Full scan settings: 70k resolution (*m/z* 200), AGC target 3e6, maximum injection time 120 ms. dd settings: minimum AGC target 8e3, intensity threshold 7.3e4, charge exclusion: unassigned, 1, 8, >8, peptide match preferred, exclude isotopes on, dynamic exclusion 45s. MS2 settings: microscans 1, resolution 35k (*m/z* 200), AGC target 2e5, maximum injection time 110 ms, isolation window 2.0 *m/z*, isolation offset 0.0 m/z, spectrum data type profile.

#### Data Analysis

The raw data was processed using Proteome Discoverer 2.4.0.305 software (Thermo Scientific) and searched against the UniProt database for the *Rattus norvegicus* Proteome 2020_02 together with a common contaminant database from MaxQuant (version 1.6.2.6, Max Planck Institute of Biochemistry, Munich, Germany). The Sequest HT search engine was used to identify tryptic peptides. The ion mass tolerance was 10 ppm for precursor ions and 0.02 Da for fragment ions. Maximum allowed missing cleavage sites was set 2. Cysteine carbamidomethylation was defined as constant modification. Methionine oxidation, protein N-terminus acetylation, Met-loss and Met-loss+acetylation, were defined as variable modifications. Peptide confidence was set to high. The processing node Percolator was enabled with the following settings: maximum delta Cn 0.05; decoy database search target FDR 1%, validation based on q-value. Protein label free quantitation was performed with the Minora feature detector node at the processing step. Precursor ions quantification was performing at the processing step with the following parameters: Peptides to use unique plus razor, precursor abundance based on intensity and normalization based on total peptide amount.

#### Protein Functional Enrichment Analysis

The enrichment analyses were performed with GSEA (Gene Set Enrichment Analysis). The biological process (BP), cellular component (CC), and molecular function (MF) were conducted based on the Gene Ontology (GO) terms of each protein. Further, Kyoto Encyclopedia of Genes and Genomes (KEGG) and Reactome analysis was employed and the pathways or GO terms with p-value 0.05 were considered significant.

## Results

### Behavioral assessment

To assess the impact of the social instability stress on the sociability of adolescent rats, in two distinct periods, we evaluated social recognition and social novelty preference in a three-chamber apparatus. In the sociability test a novel same-sex rat (stranger 1) was placed in one side chamber within a wire cup, which prevented direct social interaction while permitting transmission of olfactory, auditory and visual stimuli. An empty wire cup (novel object) was placed in the other side chamber and served as the novel nonsocial object (Fig. 2 A). The center chamber, considered neutral, contained neither objects nor animals. Regardless of sex, the control and stress groups (PND 30 and PND 45) spent more time in the chamber containing the novel animal than the non-social chambers (*p* < 0.05 by Sidak *post-hoc* tests following two-way ANOVA with the factors of condition and sex). Notably, male and female at PND 45 showed more sociability behavior than animals at PND 30 [male: Age F_(1,14)_ = 115.5; *p*<0.0001; female Age F_(1,14)_ = 142.0; *p*<0.0001] (Fig. 2 B, C). Concerning, social novelty (Fig. 2 D), the first rat (stranger 1) remained in one side chamber and was now a familiar social stimulus. A new same-sex rat (stranger 2) was introduced in the other side chamber. All groups demonstrated the preference for investigating the stranger 2 over the stranger 1 (Fig. 2 E, F). No change in social preference behavior was observed after the social stress condition in both sex or aged.

In order to understand if the SIS protocol affected social status, we evaluated social interactions in the Home Cage Change Test, both in male and female rats (Fig. 3 A). This test is a social interaction paradigm in which subtle dominance-subordination relationships were evaluated when animals re-establish their dominance-subordination relationships after a cage change in the new home-cage Magara *et al*. (2015). All behaviors were categorized into neutral, dominance, aggressive or submissive, according to Magara *et al*. (2015). The social instability stress did not change the profile of the behavioral categories in the novel cage test of male or female rats, in either age (Fig. 3 B, C). In addition, we analyzed the behaviors associated with the Dominance behavioral category, and the descriptive parameters are displayed in Figure 3 D. The duration of mount behavior was lower in the PND 30 and PND 45 male stress [male: Stress [F_(1,7)_ = 15.47; *p*=0.0057] (Fig. 3 E, G) and in the PND 30 female rats submitted to SIS than in the respective controls (*p*<0.0032) while, in the PND 45 females, there was no difference between the groups (Fig. 3 F). At this age, SIS females spent more time in anogenital sniffing (*p*=0.0393; Fig. 3 H).

**Figure 3.**
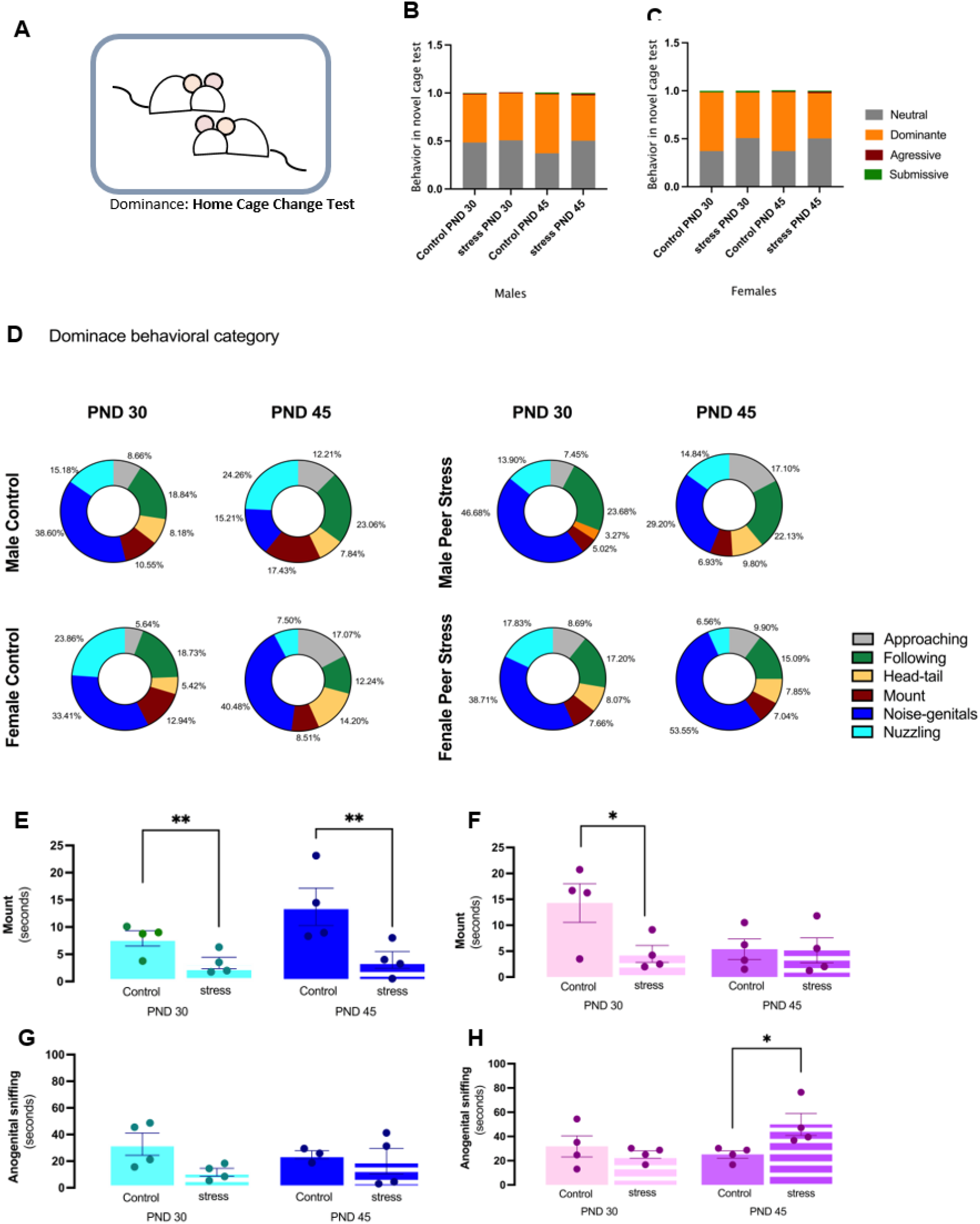
Effect of SIS on mild and late adolescence in social hierarchy assessed using the novel cage test. (**A**) Schematic representation of the Novel cage test. Duration of coping styles are expressed as fraction of the total behavior scored per animal in (**B**) male and (**C**) female rat. (**D**) Representative percentage of dominance category in control and SIS male and female rats. Data are presented as mean. (**E, F**) Duration of mount and (**G, H**) anogenital sniffing behavior mild and late male and female adolescents during the novel cage test. Two-way ANOVA with Sidak’
ss comparisons. *n*= 4 per group. Data are expressed as mean ± SEM. \**p* ≤ 0.05

### SIS affects the proteomic signature of mild, but not late, adolescent rats

In order to clarify how the SIS protocol hinders normal brain development, and whether a) there is a period of adolescence more vulnerable to alterations of the social environment and b) whether it affects males and females in the same manner, we performed a proteomics analysis of the PFC. For this, samples of three animals were used per group and alterations in the proteome were investigated via the previously established iTRAQ-based quantitative approach. In the current study, a total of 2498 distinct proteins were identified (Supplementary Table 2). The fold change values and *p*-value were calculated for all proteins and only those proteins with *p* < 0.05 (Student’
ss *t*-test) and a fold change > 1.3 were considered differentially expressed and select for further analysis.

We carried out hierarchical clustering analysis on the differentially expressed proteins (DEPs), and the result is displayed in Figure 4 A. In this study, sample information was on the x-axis, and differential proteins were on the y-axis. The color depth indicated the expression levels. Red was used to indicate upregulated proteins, and blue represented apparent downregulated proteins. Hierarchical clustering demonstrated most similarities among groups are related to age, regardless of sex or home-cage environment.

**Figure 4.**
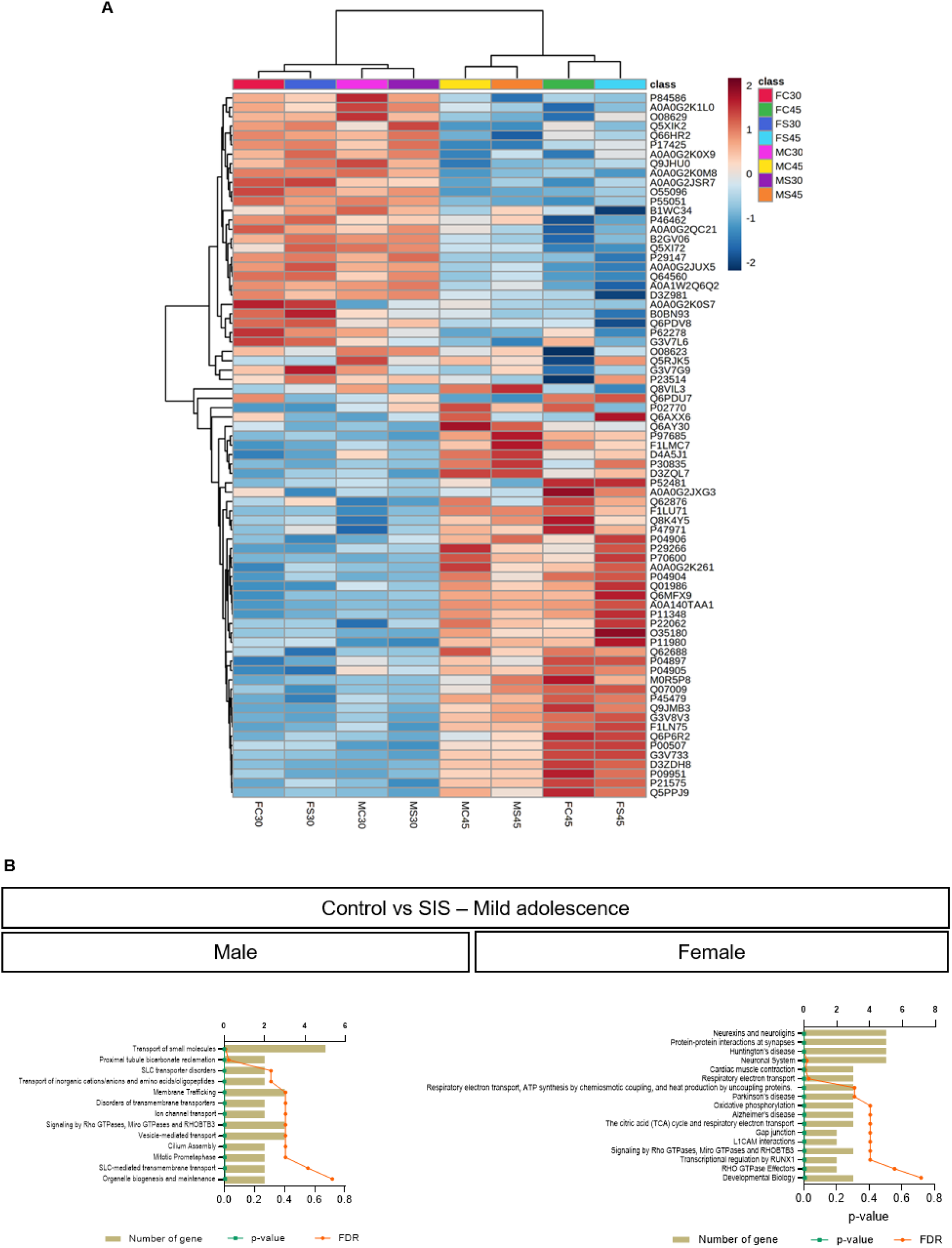
Heatmap of top 75 quantified proteins in the dataset. (**A**) Higher expressions are indicated by red and lower by green. The expression levels are indicated by different intensities of colors, which are represented by the color-key bar with a log2-scale. (**B**) The significantly enriched pathways in the PFC from male and female mild adolescents after SIS procedure.

However, differential proteomic analysis in the PFC of control vs stress age- and sex-paired groups, revealed 5 down- and 10 up-regulated proteins in the male group and 4 down- and 12 up-regulated proteins in the female group, both in mild adolescence. On the other hand, no significantly altered proteins were observed in male or females in late adolescence. The enriched GO terms of development-related proteins of rat’s PFC were predicted using GSEA-MSigDB. The 15 differentially expressed proteins from mild adolescent males were categorized according to GO enrichment analysis, and 100, 32, 19 and 12 terms were identified to be significantly enriched (*p*<0.05) in biological process (BP), cellular component (CC), molecular function (MF) and pathway, respectively (Supplementary Table 3). Similarly, the 16 differentially expressed proteins from the mild adolescence female group were categorized according to GO enrichment analysis, and 100, 48, 9 and 9 terms were identified to be significantly enriched (*p*<0.05) in BP, CC, MF and pathway (Supplementary Table 3). Upon performing the pathway analysis (considering Reactome and KEGG) the dysregulated proteins were mainly involved in transport of small molecules in SIS-male group (Fig. 4 B). In contrast, the SIS-female group showed pathways as neurexins and neuroligins, protein-protein interactions at synapses and neuronal system (Fig. 4 B). The top down-regulated proteins in the SIS-male group include Slc4a4 and Basp1 and the up-regulated are Slc25a12 and Exoc3. In the other hand, in the SIS-female group, the top down-regulated proteins include Hist2h3c2 and Ndufs7 and up-regulated Dlgap3 and Cntnap1. These data evidence the contrast of the response to stress at mild adolescence according to the sex and suggest that the molecular mechanisms underlying responses to stress are different among these groups.

### Social instability affects the normal development of adolescent rats differently on males and females

Regarding the control-SIS condition, male and female does not show any similarity between the DEPs, as we showed in Venn diagram (Fig. 5 A-C). Next we assessed how SIS affected the development of adolescent rats. For this, we compared the proteome of male PND 30 with male PND 45 and the same process was performed in the female group. We founded higher differences in the groups of animals that underwent SIS protocol compared to control animals, indicating that SIS induced alterations on the developmental process. In male control animals, comparing PND-30 and PND-45 rats, there was a total of 46 DEPs, 28 up- and 18 down-regulated (Supplementary Table 4). In the females, there was 43 DEPs, 24 up- and 19 down-regulated (Supplementary Table 4). Of these, there were 3 proteins in common between the groups – Igsf8, Tnc and Gsta3 (Supplementary Figure 2). On the other hand, comparing animals subjected to the SIS protocol, the proteomic analysis revealed 43 DEPs (15 down and 28 up regulated proteins) in the male group, 74 DEPs (49 down and 25 up regulated protein) in the female group when compared PND-30 and PND-45 adolescent rats submitted to SIS procedure (Fig. 5 A; Supplementary Table 3 E, F, respectively). Importantly, the overlay between the control and SIS animals was minimal.

**Figure 5.**
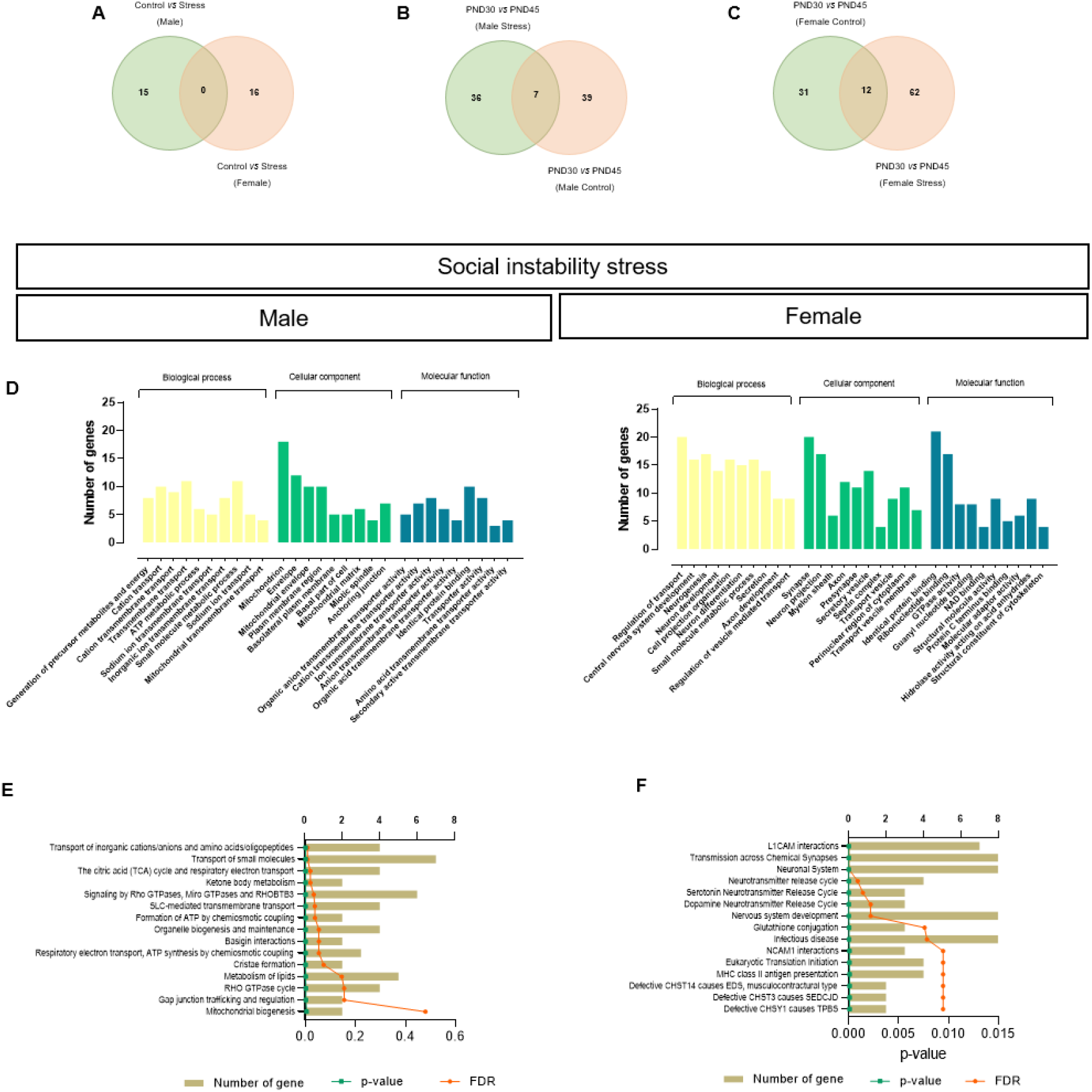
Venn diagrams comparing differentially expressed proteins on the PFC of (**A**) control females and males at PND 30 (sex-specific alterations); and mild vs late adolescence of males (**B**) and females (**C**) in control conditions or subjected to SIS (developmental alterations). The top ten enriched terms from SIS groups during development in male and female in biological process (BP), cellular component (CC), and molecular function (MF) (**D**). The significantly enriched pathways from male (**E**) and female (**F**) adolescents after SIS procedure in the PFC during development.

In Figure 5, we demonstrate 7 DEPs in common between control and SIS-subjected male rats (Fig. 5 B), while females had 12 DEPs in common (Fig. 5 C). These 12 proteins in common seem to be related with myelin sheaths, suggesting a trait that occurs during female rat development, regardless of SIS. The 43 DEPs from male group were categorized according to GO enrichment analysis, and 100, 58, 43 e 22 terms were identified to be significantly enriched (*p*<0.05) in biological process (BP), cellular component (CC), molecular function (MF) and pathway. The 74 differentially expressed proteins from female group were categorized according to GO enrichment analysis, and 100, 96, 61 e 100 terms were identified to be significantly enriched (*p*<0.05) in BP, CC, MF and pathway. The top 10 BP, CC and MF for male and female SIS-group was showed in Fig. 5 D.

Performing the pathway analysis through Reactome, the DEPs in male SIS-aged male group were generally involved in transport of small molecules, signaling by Rho GTPases, MIRO GTPases and RHOBTB3 and metabolism of lipids (Fig. 5 E). On the other hand, the SIS-aged female group exhibited L1CAM interactions, transmission across chemical synapses nervous system development and infectious diseases (Fig. 5 F). This suggested that these stressed rats had distinct molecular patterns as responsive to SIS condition, age- and gender-dependent.

## Discussion

Studies concerning adverse social experience during the adolescence period in animal models provide new insights on risk and vulnerability factors for mental health disorders at later stages of life (Jankord *et al*., 2011). Particularly, social experiences in adolescence are considered important determinants to modulate the social behavior. The variation in socio-environmental context and frequent confrontations with novel situations and emotions that occur during adolescence can be understood as expressive stressful life events (Buwalda *et al*., 2011). Hence, we conducted the social instability stress (SIS) procedure adapted from McCormick *et al* (2010) that differs from the original in three main points: 1) Animals were not previous isolated and restrained in small containers for 1 hour - this modification in protocol had as purpose eliminating the impact on animal behavior and molecular changes promoted by social isolation in animals (for review Mumtaz *et al*., 2018); 2) The procedure lasted 7 instead of 15 days; and 3) animals had two new partners each day, instead of one new partner each day. However, in both protocols all rats interacted with 15 different peers. This allowed us to take into consideration, besides sex, different stages of adolescence (mild *vs* late), but also removed the isolation and restraint factors, known stressors *per si* (for review Mumtaz *et al*., 2018). Therefore, the present study sought to examine the influence of social peer stress experiences during the development of the social brain without any other stress involved, such as isolation.

Our findings indicate that the social motivation and recognition of adolescent rats are not significantly affected by SIS (without isolation) in both male and female rats when compared with age- and sex-matched control rats. Importantly, we show that differences in social behavior in communication of dominance-subordination relationships in a Home Cage Change Test are sex and age dependent.

The ability of adolescent rats to process information used to identify and interpret social signals were evaluated as well as the behavior flexibility to express appropriate social behaviors in a new environment. We started by estimating social motivation and social recognition in the 3-Chamber Sociability and Social Novelty test in rats (Crawley, 2004). Social motivation is the essence of developing other adaptive social behaviors (Bicks *et al*., 2015), and recognizing others is a critical skill for proper social behavior (Schweinfurth, 2020). We found that the social stress experienced during mild and late adolescence did not affect sociability and social recognition in male and female of Sprague Dawley rats in the two-time points evaluated. These findings contradict others that reported that social stressors impair social recognition memory and decreased social interactions in Long Evans rats (Hodges *et al*., 2017; Marcolin *et al*., 2020; Schaack *et al*. 2021). However, Graf and colleagues (2023) found that Wistar rat exposed to SIS improved social recognition performance with 30 min but not for 90 min retention interval. Wistar rats’ strain and Sprague Dawley rats, in general, show lower stress responses which seem to be associated with lower hypothalamic-pituitary-adrenal activity when compared with other strains such as Long Evans rats (Tannahillet *et al*., 1988; Sanch’ ss-ollé *et al*., 2021). Theses strain-specific effects of social stress highlight the importance of identifying encoded genetic variants that confer resilience/vulnerability to social stress.

We also found that age, independent of SIS, is a significant factor in the expression of sociability but not in social recognition. Late adolescent rats of both sexes are more highly motivated to interact with a social stimulus than in early adolescent rats (male and females). It has been demonstrated that adolescence is a period of transitions involving increased of independence and shifts in relationships from parents to peers, and like adolescent humans, rats undergoing adolescence showed a marked increase in the amount of time spent in social investigation and interaction with peers (Spear, 2000). This higher motivation to approach, explore, and interact with others in late adolescence may have an adaptive value during this developmental period, contributing to explore and creating new relationships to form groups or to find mate.

On the other hand, age does not affect social recognition memory in both sexes. Despite very few papers reporting age effects on social memory across ages. Markham and colleagues (2007) also reported that age-related changes in social recognition memory were undetected. However, Graf and colleagues (2023) found that adolescents spent more time investigating the familiar rat compared to adult rats. Social recognition is an essential element for normal social functioning because it is critical for attachment formation, hierarchies, and other complex social behaviors (Bicks *et al*., 2015). As such it would be expected that social recognition starts early in life and is preserved during aging.

We further examined subtle dominance-subordination relationships in a Home Cage Change Test (Magara *et al*., 2015). Dominance hierarchies are particularly important in social group living animals (Cummins, 2000). Dominance-subordination relationships demand that animals recognize both their and others status within the group, communication skills, and social decision-making in order to behave appropriately (Cummins, 2000; Watanabe & Yamamoto, 2015). For this reason, hierarchy establishment can give information about complex social behaviors that require plasticity when animals are challenged in new social contexts.

In Home Cage Change Test differences between groups for all social behavior categories analyzed (Neutral, Dominance, Aggressive, Submissive) were not detected. However, we can observe that rats of all experimental groups display more behaviors included in dominant behavior category, as expected since this test evokes dominance-subordination behaviors in the re-establishment of a social hierarchy (Magara *et al*., 2015).

To further explore the social information from this dominance test, we analyzed each behavior of the dominant category and found that all rats SIS (without isolation) display less mount behavior when compared with age- and sex-matched control, except the PND 45 females. At this age, female controls already exhibit a low duration of mount behavior. However, SIS females, at PND 45, increase the anogenital sniffing. These results are exciting because they indicate that SIS alters social communication depending on sex and age however should be interpreted with caution because our sample size is small. Besides individual experience, it is known that animals can gain information about a potential threat through inference from a social context, and this capacity is critical for individuals’ ability to produce proper behaviors in response to stressful situations. This social information can be gained through a variety of species-specific signals (Monfils & Agee 2018) and may include facial expressions (Sotocinal *et al*., 2011), vocalizations (Brudzynski, 2013), odors cues (Zalaquett & Thiessen, 1991), sniffing behavior (Wesson, 2013) social contact (Jones & Monfils, 2016) including mount behavior.

Mounting is categorized as threat behavior that acts as a form of “ritualized” aggression (Lorenz, 1966) to maintain the social hierarchy (Scott, 1966). It is worth mentioning aggressive displays are relatively rare in laboratory rats (Blanchard & Blanchard 1990, 1988, 1977), especially in a docile strain of Sprague–Dawley rats. The reduced mount behavior by the majority of SIS rats suggests that SIS rats may reflect a copping strategy to deal with their submissive profile, taking a more passive and tolerable social interaction in response to the new environment. This social behavior could indicate that SIS adolescent males and early adolescent females exhibit a reactive coping style (Magara *et al*., 2015). This type of caution/defensive response was also observed by Jones and Monfils, (2016) in subordinate rats after the social transmission of fear. These authors quantify the nape contacts as indicators of dominance status and observed that subordinate rats counter attacking in response to nape contacts less than dominant rats.

On the other hand, late (PND 45) adolescent SIS females seem to have a different social strategy in new home-cage environment. Theses females increased time spent in anogenital sniffing with their cage mate. These results are in line with Rogers-Carter and colleagues (2018) that showed that females rats approached stressed familiar conspecifics which can influence the reciprocal social behaviors and decision-making to show more social exploration.

Anogenital sniffing is frequently displayed in the first minutes of investigation and tend to decrease throughout social interaction. This is an important behavior for rodents because transmits the sexual and emotional state of an animal. Kiyokawa and collegues (2006; 2013; 2017) showed that stressed animals can release alarm pheromones from the anogenital area that induce a stress response in recipients. (Kiyokawa *et al*., 2004; 2006). Recent studies have recognized a significant role of anogenital sniffing in social transmission of stress (Neff, 2018; Lee *et al*, 2021). SIS mild adolescent females seem to adopt a more active coping strategy (actively responding) than other experimental groups. From an ethological view, this gain of information from cage mates has evident adaptive value since may promote cooperation/affiliation establishment to prepare individuals for subsequent challenges (Monfils & Agee 2018). Our behavioral data suggests that SIS affects subtle dominance-subordination behaviors in a Home Cage Change Test and promotes both active and passive /defensive responses in adolescent rats in an age- and sex-dependent manner.

These behaviors are underlain by shared prefrontal circuits (Bicks *et al*., 2015). Prefrontal cortex (PFC) comprises a crucial element of the brain networks that have been implicated in the regulation of emotion, social cognition, and decision making (Beer & Bhanji, 2011; Franklin *et al*., 2017). Several studies showed the influence of eternal stimulus, as stress, depending on duration and intensity, can lead to an altered development of the PFC and its connected areas, like hippocampus and amygdala (for review Arnsten, 2009). Disturbance or failure in executive function are symptoms of several stress-related psychological disorders (e.g., schizophrenia, depression, PTSD). Social stress is particularly damaging during adolescence, as this is a period of dynamic growth and restructuring in the PFC renders this brain area sensitive to external and internal elements that may impact mental health later in life (Smith & Pollak, 2020).

Removing the isolation period of most SIS protocols described on literature (for review Koert *et al*., 2021) in this work revealed that the social stress by itself does not impact social cognition and hierarchic behavior neither in early nor mild phases of adolescent rats. Nonetheless, we found that mild adolescent rats are more vulnerable to molecular changes induced by SIS. Remarkably, during the mild adolescence, the DEPs between unstressed control and SIS-group are totally sex dependent. This suggested that these stressed male and female groups had distinct molecular patterns as responses to SIS procedure. In line with this, we found that the proteomic signature of unstressed male and female rats is vastly different depending on the stage of adolescence. Moreover, the proteome of SIS-subjected animals disrupted the sex-differences of the proteome of mild adolescents and decreased the proteomic alterations between males and females on late adolescence. Similarly, Walker and colleagues (2019) have described that social isolation in adolescent mice, a strong stressor, is capable of disrupting sex-specific transcriptional differences in the PFC.

Our study suggests that DEPs in SIS-male group (control *versus* SIS) at mild adolescence were primarily involved in the transport of small molecules pathway, membrane trafficking and signaling by Rho GTPas be, Miro GTPase and RHOBTB3. Among the DEPs in male group, we observed changes in the expression as down-regulation of Mtch2 and up-regulation of Slc25a12. MTCH2 has been characterized as a regulator of mitochondrial metabolism, motility, and calcium buffering (Ruggiero *et al*., 2017; Manjunath *et al*., 2020). Ruggiero and cols (2017), shows that Loss of MTCH2 decreases mitochondria motility and calcium handling and impairs hippocampal-dependent cognitive functions. Still, Lepagnol-Bestel and cols (2008), showed that Slc25a12 overexpression in prefrontal cortex may be involved in the pathophysiology of autism. The mitochondrial dysfunction has been associated as vulnerability factor for depression and stress-related psychopathologies (Morava & Kozicz, 2013; Petschner *et al*., 2018; Picard *et al*., 2018).

In parallel, pathway analysis in female mild adolescents revealed that the DEPs were involved in neurexins and neuroligins, protein-protein interaction at synapses and neuronal system. Among the DEPs in this group, we found a down-regulation of Dlg4, Dlgap3, Grm5, Shank2, Homer1 genes as compared with control age and sex-paired animals. The post-synaptic density protein (PSD95) is encoded by the Dlg4 gene, consists of a major synaptic protein and is an essential component involved in glutamatergic transmission, synaptic plasticity, and dendritic spine morphogenesis during neurodevelopment (Kim & Sheng, 2004). The dysfunction of PSD95 has been implicated in dual changes in social behavior age- and brain region-specific dependent (Coley & Gao, 2019, Feyder *et al*., 2010; Winkler *et al*., 2018). Also, studies have showed that Grm5 gene is relevant in sociability (Ramos-Prat *et al*., 2019; Barnes *et al*., 2015). The response to SIS in females (PND30) still revealed a mitochondrial dysfunction through the up-regulated Hist2h3c2, Ndufs7 and Cox4i1 and down-regulation of Uqcrq. It is known that mitochondrial function modulates neuroendocrine, metabolic and inflammatory response to psychological stress (Picard *et al*., 2015).

Although the SIS protocol did not reduce social interactions in male and female rats at PND 30, the proteomic signature in the PFC of these animals reveals that the negative social experience was able to promote considerable molecular changes mainly in females. It is possible to speculate that these changes observed immediately after the SIS may have a long-term behavioral impact in adulthood. For example, Hyer and colleagues (2021) have demonstrated that chronic stress during female adolescent rats, provoked impairment of cognitive flexibility on adulthood. On the other hand, Suo and colleagues (2013) demonstrated that predictable chronic stress in adolescence contributed to resilience against depression and anxiety in adulthood, and similar results were obtained by Mancini and colleagues (2021), demonstrating that the type of stress has a key role on the outcome in adults.

Considering development proteome signature in control groups between male and female rats, the minimal overlap of DEPs was observed: Igsf8, Tnc and Gsta3 (Supplementary Figure 2), being these proteins are involved in regulation of synaptic function and hippocampal organization, migration of neurons, specifically in neurogenic areas and oxidative stress regulation (Martínez-Lara *et al*., 2003; Ohtaka-Maruyama *et al*., 2016; Apóstolo *et al*., 2020). When we observed the interactions of proteins under the aspect of development is safe to assume that SIS affects a distinct group of proteins at the different ages analyzed. Only 7 proteins are related to development in males Tppp and Dgkh as up-regulated and Mapre1, Oxct1, Dpysl3, Tnc as down-regulated (Supplementary Table 2). Still, when we evaluate the female group we found 12 identical proteins shared in PND 30 and PND 45: Qdpr Gsta3 Hcn2 Sept4 Mog Tubb4a Gfap Syn2 as up-regulated and Cdh13 Bdh1 Dclk1 as down-regulated proteins (Supplementary Table 2). The small overlap between SIS-subjected rats and control animals demonstrates that SIS induced a reprogramming of the developmental proteome.

Given the fact that the altered proteins are different in distinct age groups, we performed the analysis considering only the condition of social stress at different ages (PND30 *versus* PND45 SIS-groups). The down-regulated pathway analysis in SIS male group are mainly involved with metabolism and whereas the up-regulated pathway was related to Signaling by Rho GTPase, Miro GTPase and RHOBTB3. Considering the highlight to mitochondria and plasm membrane as cellular component in GO analysis, Miro, atypical Rho GTPase protein located on the mitochondrial outer membrane, plays a critical role in axonal transport, dynamics process and homeostasis of mitochondria supporting the neuronal energy demand and neuronal survival, so dysfunction of this proteins plays a key role in several neurodegenerative diseases by impairing mitochondrial functions (Calkins *et al*., 2011; Birsa *et al*., 2013; Panchal *et al*., 2021).

The GO analysis to cellular component revealed the involvement of synapse, neuron projection and myelin sheath in the female SIS-group that culminate in up-regulation of neuronal system and transmission across chemical synapses and down-regulation of translation and activation of the mRNA upon binding of the cap-binding complex and eIFs, and subsequent binding to 43S. Our data showed the dysregulated expression of translation initiation factors EIF2S2, EIF4H and EIF3J. The initiation of protein synthesis is suppressed under several stress conditions, inducing phosphorylation of the α-subunit of the eukaryotic initiation factor 2 (eIF2α). Studies have indicated that α-subunit of the eukaryotic initiation factor 2 (eIF2α)-mediated translational control regulates synaptic plasticity (Costa-Mattioli *et al*., 2009; Bellato *et al*., 2016). Furthermore, control of translation is fundamental for fine-tuning of gene expression and plays a critical role in development, cellular growth, proliferation, differentiation, synaptic plasticity, memory and learning (Sonenberg & Hinnebusch, 2009). Recent studies have suggested a sex difference in synaptic pruning of the PFC during adolescence. Female rats appear to have an earlier peak in synapses than males which could alter the window of sensitivity to exposures between the sexes (Drzewiecki *et al*., 2016; Page & Coutellier, 2018).

The main limitation of this study was the small sample size (Novel Home Cage test and PFC sample n=3/each group). In the future, the proteomic modifications in the prefrontal cortex of male and female rats with increased sample size should be performed. In addition, behavioral studies in adult animals are required to elucidate the impact of early life social instability stress.

## Conclusion

We found that peer-insatiability stress (without 1 hour of isolation) does not affect social recognition and motivation but leads to distinct coping strategies for dominance-subordination relationships, with peers in a new environment, depending on age and sex. SIS adolescent males and early adolescent females display a more defensive coping strategy, whereas SIS at late adolescent females showed a more active coping strategy. The social communication involved in the hierarchy establishment can give information about complex social behaviors, especially after stressful events, which are particularly important in social group-living animals.

Moreover, SIS also modifies the proteomic signature of females, affecting the expression of proteins and pathways which are involved in increased vulnerability to mental disorders. SIS leads to subtle modifications at the molecular level that hamper normal development. Proteomics analysis of the PFC revealed the molecular modifications, identifying biomarkers, and elucidating the mechanisms of social instability stress and the importance of sex and age differences. The impact of developmental psychosocial stress on the circuitry of the PFC and resultant phenotypes must be taken into account with a focus on the importance of considering sex differences to build a better understanding of developmental influences on adult disorders.

Collectively, these findings suggest that at different time points, adolescent social insatiability stress can distinctively affect social responses to dominance-subordination relationships, and may act as a priming event that impacts the expression of proteins and pathways that can lead to long-term neuropsychiatric disorders, in a sex-dependent manner. Further studies are needed to address the impact of these alterations in adulthood.

## Acknowledgments

This work was financed through the FEDER-Fundo Europeu de Desenvolvimento Regional funds through the COMPETE 2020 - Operational Programme for Competitiveness and Internationalisation (POCI), Portugal 2020, and FCT - Fundação para a Ciência e a Tecnologia/Ministério da Ciência, Tecnologia e Ensino Superior in the framework of the project POCI-01-0145-FEDER-032231 (PTDC/SAU-TOX/32231/2017); and FCT and Orçamento do Estado in the framework of the project AMAGALHÃES - FCT_FProgramático_Neuro /i3S/1701/2022. RLA was supported by an FCT grant (PD/BD/114266/2016).

The authors acknowledge the support of the following i3S Scientific Platforms: Animal Facility and Proteomics.

