## Supplementary Table 1 for "Prefrontal cortex proteomic alteration after social instability stress in adolescents rats": Supplementary Table 1.Ethograms of behaviors in the novel cage tests.pdf

Table 1. Ethograms of behaviors in the novel cage tests (Magara *et al*., 2015)

.

| Category | Behavior | Description |
|---|---|---|
| **Neutral behavior** | | |
| | *Head-head* | The head of the rat touches the head of the other rat |
| | *Nose-side* | The rat sniffs between the ventral region and the back of the other rat |
| | *Nose-nose* | The rat sniffs the other rat´s nose in an equal sniff |
| | *Passing* | The rat passes the other rat either in a direct meeting or from behind |
| **Dominante behavior** | | |
| | *Head-tail* | The head of the rat touches the tail of the other rat |
| | *Nose-genitals* | The nose of the rat touches the genitals of the other rat |
| | *Folllowing* | The rat follows the other rat for more than two steps |
| | *Approaching* | The rat is walking or running more than three steps without hesitation |
| | *Nuzzling* | The rat sniffs/bites/grooms the other rat in the area between the tip of the nose and the ventral region |
| | *Mount 1* | The rat rears and leans its front legs on the other rat´s back from behind |
| **Aggressive behavior** | | |
| | *Mount 2* | The rat rears and leans its front legs on the other rat´s back from behind and makes copulation movements |
| | *Chasing* | The rat runs after the other rat for more than two step |
| | *Fight* | Very rapid rolling, jumping and biting of both animas while begin in close contact |
| **Submissive behavior** | *Avoiding* | The rat moves or faces in a direction away from the other rat when the other rat is approaching |
| | *Crowing under* | The rat is crawling under the other rat |
| | *Submissive posture* | The rat is lying on its back with the other rat standing and/or leaning over its ventral part |
